# HARMONIES: A Hybrid Approach for Microbiome Networks Inference via Exploiting Sparsity

**DOI:** 10.1101/2020.03.16.993857

**Authors:** Shuang Jiang, Guanghua Xiao, Andrew Young Koh, Bo Yao, Qiwei Li, Xiaowei Zhan

**Affiliations:** Department of Statistical Sciences, Southern Methodist University, Dallas, TX, USA; Quantitative Biomedical Research Center, University of Texas Southwestern Medical Center, Dallas, TX, USA; Departments of Pediatrics, Departments of Microbiology, University of Texas Southwestern Medical Center, Dallas, TX, USA; Department of Mathematical Sciences, The University of Texas at Dallas, Richardson, TX, USA

## Abstract

The human microbiome is a collection of microorganisms. They form complex communities and collectively affect host health. Recently, the advances in next-generation sequencing technology enable the high-throughput profiling of the human microbiome. This calls for a statistical model to construct microbial networks from the microbiome sequencing count data. As microbiome count data are high-dimensional and suffer from uneven sampling depth, over-dispersion, and zero-inflation, these characteristics can bias the network estimation and require specialized analytical tools. Here we propose a general framework, HARMONIES, a Hybrid Approach foR MicrobiOme Network Inferences via Exploiting Sparsity, to infer a sparse microbiome network. HARMONIES first utilizes a zero-inflated negative binomial (ZINB) distribution to model the skewness and excess zeros in the microbiome data, as well as incorporates a stochastic process prior for sample-wise normalization. This approach infers a sparse and stable network by imposing non-trivial regularizations based on the Gaussian graphical model. In comprehensive simulation studies, HARMONIES outperformed four other commonly used methods. When using published microbiome data from a colorectal cancer study, it discovered a novel community with disease-enriched bacteria. In summary, HARMONIES is a novel and useful statistical framework for microbiome network inference, and it is available at https://github.com/shuangj00/HARMONIES.

## 1 Introduction

Microbiota form complex community structures and collectively affect human health. Studying their relationship as a network can provide key insights into their biological mechanisms. The exponentially growing large datasets made available by next-generation sequencing (NGS) technology Metzker (2010), such as 16S rRNA gene and metagenomic profiling, motivate the development of statistic tools to quantitatively study the microbial organisms. While the number of discovered microbial taxa continues to increase, our knowledge of their interactive relationships is severely lacking. Understanding the structural organization of the human microbiome plays a vital role in revealing how the microbial taxa are collaborating or competing with each other under different physiologic conditions.

In sequencing-based microbial association studies, the enormous amount of NGS data can be summarized in a sample-by-taxon count table where each entry is a proxy to the underlying true abundance. However, there is no simple relationship between the true abundances and the observed counts. Additionally, microbiome sequencing data usually have an inflated amount of zeros, uneven sequencing depths across samples, and over-dispersion. Initial attempts of constructing microbial association networks with this type of data, Ban et al. (2015); Lo and Marculescu (2017) first transformed the microbiome sequencing counts into their compositional formula. Specifically, a count was normalized to its proportion in the respective sample. Then, each sample were transformed by a choice of log-ratio transformations to remove the unit-sum constraint of the compositional data. While this type of normalization is simple to implement and preserves the original ordering of the counts in a sample, it fails to capture the sample to sample variation and it overlooks the excess zeros in the microbiome data. Note that these zeros can be attributed to biological or technical reasons: either certain taxa are not present among samples, or they are not sequenced due to insufficient sequencing depths. As the existing logarithmic transformation neglects the difference between these two types of zeros, it can lead to a biased estimation of the network structure. Thus, we propose a model-based normalization strategy for microbiome count data. Our normalization method simultaneously accounts for uneven sequencing depth, zero-inflation, over-dispersion, as well as the two types of zeros. Then we use the normalized abundances to estimate microbial abundance networks.

There are two major categories of statistical methods that are often used to infer microbial abundance networks. The first type is based on a taxa abundance covariance structure. For example, Faust and Raes (2016) and Weiss et al. (2016) used pairwise Pearson correlations to represent edge weights. This simple inference could be problematic since two variables (i.e. taxa) may be connected in the network due to their confounding variables (Gevers et al., 2014). The other type aims to estimate taxa abundance partial correlations, removing confounding effects. Kurtz et al. (2015) proposed a statistical model for inferring microbial ecological network, which is based on estimating the precision matrix (via exploiting sparsity) of a Gaussian multivariate model and relies on graphical lasso (Glasso) Friedman et al. (2008). However, their data normalization step needs to be improved to account for unique characteristics observed in microbiome count data.

In this paper, we propose a general framework, HARMONIES (a Hybrid Approach foR MicrobiOme Network Inferences via Exploiting Sparsity), to infer the microbiome networks. It consists of two major steps: (1) normalization of the microbiome count data by fitting a zero-inflated negative binomial (ZINB) model with the Dirichlet process prior (DPP), (2) application of Glasso to ensure sparsity and using a stability-based approach to select the tuning parameter in Glasso. The estimated network contains the information of both the degree and the direction of associations between taxa, which facilitates the biological interpretation. We demonstrated that HARMONIES could outperform other state-of-the-art tools on extensive simulated and synthetic data. Further, we used HARMONIES to uncover unique associations between disease-specific genera from microbiome profiling data generated from a colorectal cancer study. Based on these results, HARMONIES will be a valuable statistical model to understand the complex microbial associations in microbiome studies. The `R` package `HARMONIES` is freely available at https://github.com/shuangj00/HARMONIES.

## 2 Methods

### 2.1 Microbiome count data normalization

Let ***Y*** denote the *n*-by-*p* taxonomic count matrix obtained from either the 16S rRNA or the metagenomic shotgun sequencing (MSS) technology. Each entry *y*_*ij*_, *i* = 1,…,*n, j* = 1,…,*p* is a non-negative integer, indicating the total reads related to taxon *j* observed in sample *i*. It is recommended that all chosen taxa should be at the same taxonomic level (e.g., OTU for 16S rRNA or species for MSS) in that mixing different taxonomic levels in the proposed model could lead to improper biological interpretation. As the real microbiome data are characterized by zero-inflation and over-dispersion, we model *y*_*ij*_ through a zero-inflated negative binomial (ZINB) model as

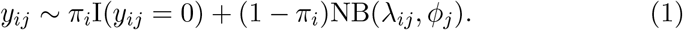

The first component in the Equation (1) models whether zeros come from a degenerate distribution with a point mass at zero. It can be interpreted as the “extra” zeros due to insufficient sequencing effort. We can assume there exists a true underlying abundance for the taxon in its sample, but we fail to observe it with the mixture probability *π*_*i*_ representing the proportion of “extra” zeros in sample *i*. The second component, NB(*λ*_*ij*_, *ϕ*_*j*_), models the “true” zeros and all the nonzero observed counts. i.e., counts generated from a negative binomial (NB) distribution with the expectation of *λ*_*ij*_ and dispersion 1/*ϕ*_*j*_. Here, “true” zero refers to a taxon that is truly absent in the corresponding sample. The variance of the random variable from NB distribution, under the current parameterization equals to 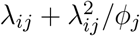 Smaller values of *ϕ*_*j*_ can lead to over-dispersion.

To avoid explicitly fixing the value of *π*_*i*_’s and *ϕ*_*j*_’s, we use a Bayesian hierarchical model for parameter inference. First, we rewrite the model (1) by introducing a binary indicator variable *η_ij_ ~* Bernoulli(*π*_*i*_), such that *y*_*ij*_ = 0 if *η*_*ij*_ = 1, and *y*_*ij*_ ~ *NB*(*λ*_*ij*_, *ϕ*_*j*_) if *η*_*ij*_ = 0. Then, we formulate a beta-Bernoulli prior of *η*_*ij*_ by assuming *π_i_ ~* Beta(*a*_*π*_, *b*_*π*_), and we let *a*_*π*_ = *b*_*π*_ = 1 to obtain a non-informative prior on *η_ij_*. We specify independent Gamma prior Ga(*a_ϕ_, b_ϕ_*) for each dispersion parameter *ϕ_j_*. Letting *a_ϕ_* = *b_ϕ_* = 0.001 results in a weakly informative gamma prior.

The mean parameter of the NB distribution, *λ_ij_*, contains the key information of the true underlying abundance of the corresponding count. As *λ_ij_* is affected by the varying sequencing effort across samples, we use a multiplicative characterization of the NB mean to justify the latent heterogeneity in microbiome sequencing data. Specifically, we assume *λ_ij_* = *s_i_α_ij_*. Here, *s_i_* is the sample-specific size factor that captures the variation in sequencing depth across samples, and *α_ij_* is the normalized abundance of taxon *j* in sample *i*.

In parameter estimation, one need to ensure identifiability between *s_i_* and *α_ij_*. For example, *s_i_* can be the reciprocal of the total number of reads in sample *i*. The resulted *α_ij_* is often called relative abundance, which represents the proportion of taxon *j* in sample *i*. In this setting, the relative abundances of all the taxa in one sample always sum up to 1. Similarly, other methods have been proposed with different constraints for normalizing the sequencing data Robinson and Oshlack (2010); Anders and Huber (2010); Bullard et al. (2010); Paulson et al. (2013). Some normalization methods can perform better than the others in the downstream analysis (e.g., the differential abundance analysis) under certain settings. From a Bayesian perspective, fixing the values of *s_i_*’s imposes a strongly informative prior in model inference. Hence, all these methods could bias the estimations of other model parameters and degrade the performance of downstream analyses. We thus propose a regularizing prior with a stochastic constraint for estimating *s_i_*’s. Our method can simultaneously infer the size factor and other model parameters. In particular, we adopt the following mixture model for *s_i_*,

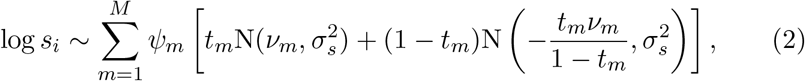

where *ψ_m_* is the weight for outer mixtures of the *m*th component. The inner mixture of the *m*th component consists of two Gaussian distributions with *t_m_* and 1 *− t_m_* as weights, respectively. It is straightforward to see that the inner mixture has a mean of zero and thus ensuring the stochastic constraint of E(log *s_i_*) = 0. For the outer mixtures, *M* is an arbitrary large positive integer. Letting *M → ∞* and defining the weight *ψ_m_* by the stick-breaking procedure (i.e., *ψ*_1_ = *V*_1_, 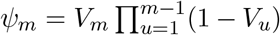), *m* = 1, 2, …) makes model (2) a special case of Dirichlet process mixture models. This class of Bayesian nonparametric infinite mixtures is widely used in quantifying the model uncertainty and allowing for flexibility in parameter estimation Kyung et al. (2011); Taddy et al. (2012). In particular, this Dirichlet process prior (DPP) has been used to account for sample heterogeneity since it is able to capture multi-modality and skewness in a distribution Li et al. (2017); Lee and Sison-Mangus (2018). In practice, we set *M* to be a large positive integer, and adopt the following hyper-prior distributions for the parameters in (2) such that *ν_m_ ~* N(0*, τ_ν_*)*, t_m_ ~* Beta(*a_t_, b_t_*), and *V_m_ ~* Beta(*a_m_, b_m_*). We further set 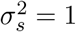 to complete the parameter specification in the DPP prior.

In our model, the normalized abundance matrix ***A*** = {*α_ij_*} represents the true underlying abundance of the original count matrix. We further assume 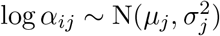. This variance-stabilizing transformation on each *α_ij_* not only reduces the skewness of the normalized abundance, but converts the nonnegative *α_ij_* to a real number. We apply the following conjugate setting to specify the priors for *μ*_*j*_ and 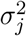, *j* = 1,…,*p*. We let 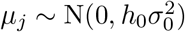 and 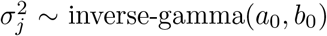. After integrating out *μ*_*j*_ and 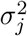, the prior of the normalized abundances of taxon *j* follows a non-standardized Student’s t-distribution, i.e.,

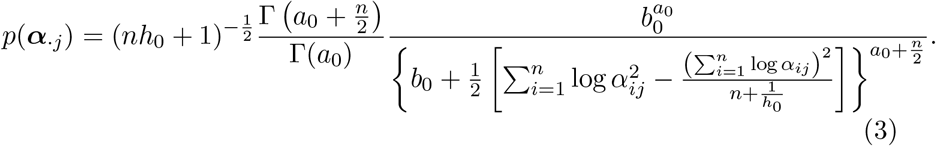

As for the fixed parameters *a*_0_*, b*_0_*, h*_0_ and 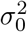, we follow Li et al. (2019) and set *a*_0_ = 2, *b*_0_ = 1 to obtain a weakly informative prior for 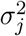. We fix 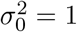 and let *h*_0_ = 10 such that the normal prior on *μ_j_* is fairly flat. We adopt the following prior specification for the rest model parameters. First, we assume an noninformative prior for each *π_i_* by letting *a_π_* = *b_π_* = 1. Next, we specify *a_ϕ_* = *b_ϕ_* = 0.001 in the Gamma prior distribution for all *ϕ_j_*’s. Then, we apply the following prior setting for the DPP: *M* = *n*/2, *σ*_*s*_ = 1, *τ*_*ν*_ = 1, *a*_*t*_ = *b*_*t*_ = 1, and *a*_*m*_ = *b*_*m*_ = 1.

The logarithmic scale of ***A***, denoted as ***Z*** = log ***A***, represents the normalized microbiome abundances on the log scale. We use Markov chain Monte Carlo (MCMC) algorithm for model parameter estimation (see details in the supplementary material), and calculate the posterior mean of ***Z*** to fit the Gaussian graphical model in the next step. Since the observed zero counts may not always represent the absence of taxa in the samples, we treat these zeros differently in the matrix ***Z***. We categorize the two types of zeros (“extra” and “true” zeros) based on the estimated *η_ij_* for each observed *y_ij_* = 0 in the data. In particular, suppose that we observe *L* zeros in total. We calculate the marginal posterior probability of being 1 for each *η_l_, l* = 1,…,*L* as 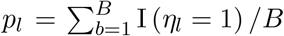, where I(·) is the indicator function, and *B* is the number of MCMC iteration after burn-in. This marginal posterior probability *p_l_* represents the proportion of MCMC iterations in which the *l*th 0 is essentially a missing value rather than the lowest count in the corresponding sample. Then, the observed zeros can be dichotomized by thresholding the *L* probabilities. The zeros with *p_l_* greater than the threshold are considered as “true” zeros in the data, whereas the rest are imputed by the corresponding posterior mean of log ***α_·j_***. We used the method proposed by Newton et al. (2004) to determine the threshold that controls the Bayesian false discovery rate (FDR) to be smaller than *c_η_*. Specifically, we first specify a small number *c_η_*, which is analogue to the significance level in the frequentist setting. Then we compute the threshold following Equation (4), which guarantees the imputed zeros have a Bayesian FDR to be smaller than *c*_*η*_,

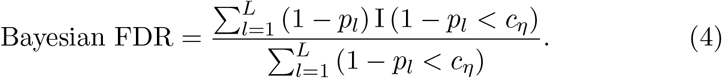

In practice, a choice of *c_η_* = 0.01 guarantees that the Bayesian FDR to be at most 0.01. We set *c_η_* = 0.05 for the simulation study and *c_η_* = 0.01 for the real data analysis.

### 2.2 Graphical model for inferring taxa-taxa association

Based on the normalized microbial abundances, we estimate their partial correlation matrix in order to construct the microbiome network under the Gaussian graphical model (GGM) framework. An undirected graph *G* = (*V, E*) is used to illustrate the associations among vertices *V* = {1,…,*p*}, representing the *p* microbial taxa. *E* = {*e_mk_*} is the collection of (undirected) edges, which is equivalently represented via a *p*-by-*p* adjacency matrix with *e_mk_* = 1 or 0 according to whether vertices *m* and *k* are directly connected in *G* or not. GGM assumes that the joint distribution of *p* vertices is multivariate Gaussian N(***μ***, **Σ**), yielding the following relationship between the dependency structure and the network: a zero entry in the precision matrix **Ω** = **Σ**^*−*1^ indicates the corresponding vertices are conditional independent, and there is no edge between them in graph *G*. Hence, a GGM can be defined in terms of the pairwise conditional independence. If ***X*** ~ N(***μ***, **Ω**), then

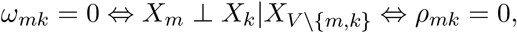

where 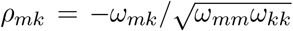 is the partial correlation between vertices *m* and *k*, representing the degree and direction of association between two vertices, conditional on the rest variables. Consequently, learning the network is equivalent to estimating the precision matrix **Ω**. For real microbiome data, we set the taxa (on the same taxonomic level) as vertices. Hence, a zero partial correlation in the precision matrix can be interpreted as no association between the corresponding pair of taxa, while a nonzero partial correlation can be interpreted as cooperative or competing associations between that taxa pair.

In biological applications, we often require a sparse and stable estimation of the precision matrix **Ω**. The sparsity can be achieved by imposing *l*_1_-penalized log-likelihood,

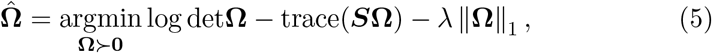

where ***S*** is the sample covariance matrix. The coordinate descent algorithm can iteratively solve *p*. The estimated precision matrix is sparsistent (i.e., all the parameters that are zeros would be estimated as zero with probability one) Lam and Fan (2009), as Glasso theoretically guarantees a consistent recovery of the sparse graph for the *p* vertices. When *p* >> *n*, the computational efficiency is often satisfactory, and thus Glasso is widely used in studying large-scale biological networks Menéndez et al. (2010); Oh and Deasy (2014); Zhao and Duan (2019). We employ a stability-based approach to select the tuning parameter in the Glasso, which is named Stability Approach to Regularization Selection (StARS) Liu et al. (2010). This method is an improved algorithm for estimating the tuning parameter *λ* in (5). The StARS selects the optimal sparsity parameter according to the graph reproducibility under the subsampling of the original data. In general, for each *λ* along the sparsity parameter path, we first obtain random subsamples from the original data. Then we estimate the graph for each subsample using the Glasso. Next, for each sparsity parameter, we calculate the overall edge selection instability from all the graphs constructed by the subsamples. Finally, the optimal sparsity parameter *λ^*^* is chosen such that it corresponds to the smallest amount of regularization and still results in a graph instability to be lower than the pre-specified tolerance level. Liu et al. (2010) showed that StARS could provide the “sparsistent” network estimation that includes all the true associations with probability one. Further, the StARS has been widely used in biological network studies Kurtz et al. (2015); Tipton et al. (2018); Zhao and Duan (2019). Due to its excellent performance, here we adopt the StARS to select the tuning parameter for Glasso. In summary, we use the normalized abundances (on the log scale) as inputs, calculate the sparse estimation of the precision matrix using the Glasso, and use the StARS method to select *λ* in problem (5) to obtain the estimated graph that represents the microbiome network.

### 2.3 Simulation scenarios

We compare the performance of the HARMONIES and several widely used methods for inferring microbiome networks. These methods include SPIEC-EASI Kurtz et al. (2015), CClasso Fang et al. (2015) and correlation-based network estimation used in Faust and Raes (2016); Weiss et al. (2016). While the proposed model and SPIEC-EASI infer the network structure from sparse precision matrices, CClasso, and the correlation-based method utilize sparse correlation matrices to represent the network. We generated both simulated and synthetic datasets that mimic the real microbiome sequencing count data. We use ***Y***_*n*×*p*_ to denote the generated count matrix. For a comprehensive comparison, we varied the sample size and the number of taxa as *n* ∈ {60, 100, 200, 500}, and the number of taxa *p* ∈ {40, 60}.

#### 2.3.1 Generating simulated data

We generated the simulated datasets from a Dirichlet-multinomial (DM) model using the following steps: (1) to generate the binary adjacency matrix; (2) to simulate the precision matrix and the corresponding covariance matrix; (3) to generate *n* multivariate Gaussian variables based on the covariance matrix to represent the true *n* × *p* underlying taxonomic abundances, denoted as ***D***; (4) to simulate the count table ***Y***_*n* × *p*_ from a DM model, with its parameters being exp(***D***); (5) to mimic the zero-inflation in real microbiome data by randomly setting part of entries in the count table to zeros. Note that the data generative scheme is different from the model assumption, which is given in Equation (1). The detailed generative models are described below.

We began with simulating a *p*-by-*p* adjacency matrix for the *p* taxa in the network. Here, the adjacency matrix was generated according to an Erdős–Rényi (ER) model. An ER model ER(*p, ρ*) generates each edge in a graph *G* with probability *ρ* independently from every other edge. Therefore, all graphs with *p* nodes and *M* edges have equal probability of 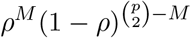. All the edges in graph *G* correspond to the 1’s in the resulted binary adjacency matrix. Next, we simulated the precision matrix **Ω** following Peng et al. (2009). We started by setting all the diagonal elements of **Ω** to be 1. Then, for the rest elements that correspond to the 1s in the adjacency matrix, we sampled their values independently from a uniform distribution Unif([−0.1, 0] ∪ [0, 0.1]). To ensure positive definiteness of the precision matrix, we followed Peng et al. (2009) by dividing each off-diagonal element by 1.5 times the sum of the absolute value of all the elements in its row. Finally, we averaged the rescaled precision matrix with its transpose and set the diagonal elements to 1. This process ensured the preceding matrix was positive definite and symmetric. The corresponding covariance matrix was set as **Σ**= **Ω**^*−*1^.

Next, we simulated *n* multivariate Gaussian variables from MN(***μ***, **Σ**) to represent the true underlying abundances ***D***. To obtain a count matrix that fully mimics the microbiome sequencing data, we generated counts from a DM model with parameter exp(***D***). Specifically, we first sampled the underlying fractional abundances for the *i*th sample from a Dirichlet distribution. The *i*th underlying fractional abundances was then denoted as ***ψ***_i_ ~ Dirichlet(exp(***D***_*i*·_)). Next, the counts in the *i*th sample were generated from Multinomial(*N_i_, **ψ**_i_*). Finally, we randomly selected *π*_0_% out of *n* × *p* counts and set them to zeros to mimic the zero-inflation observed in the real microbiome data. In general, the generative process had different assumptions from the proposed method. Under the appropriate choice of parameters, the simulated count data was zero-inflated, overdispersed, and the total reads varied largely between samples. In practice, we let *ρ* = 0.1 in the ER model. The mean parameter ***μ*** of the underlying multivariate Gaussian variable was randomly sampled from a uniform distribution Unif[0, 10]. The number of total counts across samples *N_i_, i* = 1,…,*n* was sampled from a discrete uniform distribution with range [50, 000, 100, 000]. Under each combination of *n*, *p*, and *π*_0_, we generated 50 replicated datasets by repeating the process above.

#### 2.3.2 Generating synthetic data

We generated synthetic data following the Normal-to-Anything (NorTA) approach proposed in Kurtz et al. (2015). NorTA was designed to generate multivariate random variables with an arbitrary marginal distribution from a pre-specified correlation structure (Cario and Nelson, 1997). Given the observations of *p* taxa from a real microbiome dataset, the NorTA generates the synthetic data with *n* samples as follows: (1) to calculate the *p*-by-*p* covariance matrix **Σ**_0_ from the input real dataset; (2) to generate an *n*-by-*p* matrix, denoted by ***Z***_0_, from a multivariate Gaussian distribution with mean of **0**_1×*p*_ and the covariance matrix of **Σ**_0_; (3) to use standard normal cumulative distribution function to scale values in each column of ***Z***_0_ within [0, 1]; (4) to apply the quantile function of a ZINB distribution to generate count data from those scaled values in each column of ***Z***_0_. In practice, we used `R` package `SPIEC-EASI` to implement the above data generative scheme, where the real data were from those healthy control subjects in our case study presented in Section 3.2. Under each combination of *n* and *p*, we generated 50 replicated datasets.

### 2.4 Model performance

#### 2.4.1 Alternative methods in network learning

We considered the four commonly used network learning methods. The first two methods, SPIEC-EASI-Glasso and SPIEC-EASI-mb, use the transformed microbiome abundances which are different from the normalized abundances estimated by HARMONIES. Both infer the microbial network by estimating a sparse precision matrix. The former method (SPIEC-EASI-Glasso) measures the dependency among microbiota by their partial correlation coefficients, and the latter method (SPIEC-EASI-mb) uses the “neighborhood selection” introduced by Meinshausen et al. (2006) to construct the network. The third method, denoted as Pearson-corr, calculates Pearson’s correlation coefficients between all pairs of taxa. In its estimated network, the edges correspond to large correlation coefficients. To avoid arbitrarily thresholding the correlation coefficients, the fourth method, CClasso Fang et al. (2015), directly infers a sparse correlation matrix with *l*_1_ regularization. However, as discussed in Section 1, representing the dependency structure by the correlation matrix may lead to the detection of spurious associations.

#### 2.4.2 Evaluation criteria

We quantified the model performances on the simulated data by computing their receiver operating characteristic (ROC) curves and area under the ROC curve (AUC). For the HARMONIES or SPIEC-EASI, the network inference was based on the precision matrix. Hence, under each tuning parameter of Glasso, we calculated the number of edges being true positive (TP) by directly comparing the estimated precision matrix against the true one. More specifically, we considered an edge between taxon *m* and taxon *k* to be true positive if *ω*_*mk*_ ≠ 0, 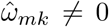, and 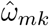 shared the same sign with *ω_mk_*. We calculated the number of true negative (TN), false positive (FP), and false negative (FN) in a similar manner. Therefore, each tuning parameter defined a point on an ROC curve. As for the correlation-based methods, we started with ranking the absolute values in the estimated correlation matrices, denoted as 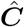. Next, we used each value as a threshold and set all the entries in 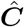 having their absolute values smaller than the current threshold to be zeros. Then, the number of TP, TN, FP, or FN was obtained by comparingthe sparse 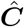 against the true partial correlation matrix. Therefore, each unique absolute value in the original estimated correlation matrix defined a point on the ROC curve.

We further used the Matthew’s correlation coefficient (MCC) to evaluate results from the simulated data. The MCC is defined as

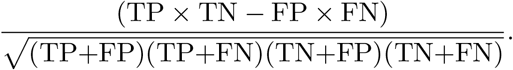

Here, the MCC was particularly suitable for evaluating network models. As the number of conditionally independent taxa pairs was assumed to be much greater than the number of dependent pairs in a sparse network, MCC was preferable to quantify the performances under such an imbalanced situation. Note that MCC ranges from [*−*1, 1], with a value close to 1 suggesting a better performance. Since each value of MCC was calculated using a given set of TP, TN, FP, and FN, we adopted the optimal choice of tuning parameter for the HARMONIES or SPIEC-EASI (with either Glasso or MB for network inference), given by StARS. As for the correlation-based methods, CClasso outputted a sparse correlation matrix. We used the result to calculate TP, TN, FP, and FN directly. For Pearson-corr, we set the threshold such that the resulted number of nonzero entries in the sparse correlation matrix was the same as the number of nonzero entries in the true sparse partial correlation matrix. In fact, this choice could favor the performance of Pearson-corr for larger sample size, as shown in Section 3.1.

To assess model performances on the synthetic datasets, we followed Kurtz et al. (2015) to use a metric called area under the precision-recall curves (AUPR), in addition to AUC. Briefly speaking, the AUPR and AUC were calculated as follows: (1) to rank all possible edges according to their confidence values; (2) to generate the precision-recall curve and the ROC curve by comparing edge inclusions against the true sparse precision matrix; (3) to calculate the area under the precision-recall curve or the ROC curve. Note that the confidence values were chosen as the edge stabilities under the optimal choice of the tuning parameter selected by StARS for HARMONIES, SPIEC-EASI-Glasso, and SPIEC-EASI-mb, while for CClasso and Pearsoncorr, *p*-values were used.

## 3 Results

### 3.1 Simulation results

Figure 1 and 2 compare the AUCs and MCCs on the simulated data under various scenarios, including varying sample sizes (*n* = 60, 100, 200, or 500), total numbers of taxa (*p* = 40 or 60), extra percentages of zeros added (*π*_0_ = 10%, or 20%). In each subfigure, the HARMONIES outperformed the alternative methods in terms of both AUC and MCC, and it maintained this advantage even with the number of sample size greatly increases. Further, a smaller sample size, a larger proportion of extra zeros added (*π*_0_ = 20%), as well as a larger number of taxa in the network (*p* = 60), would hamper the performance of all the methods, as we expected. Two modes of SPIEC-EASI, SPIEC-EASI-Glasso, and SPIEC-EASI-mb, showed very similar performances under all the scenarios, with SPIEC-EASI-Glasso having only a marginal advantage over the other. Further, we observed that the Pearson-corr method yielded higher AUCs even than the precision matrix based methods, especially when there was a lager proportion of extra zeros or larger number of taxa in the network. This result suggested that the Pearson-corr could capture the overall rank of the signal strength in the actual network. However, under a fixed cut-off value that gave a sparse correlation network, the MCCs from the Pearson-corr were always smaller than the precision matrix based methods. Note that the cut-off value we specified for Pearson’s correlation method indeed favored its performance. In general, the alternative methods considered here were able to reflect the overall rank of the signal strength by showing reasonable AUCs. However, they failed to give an accurate estimation of the network under a fixed cut-off value.

**Figure 1:**
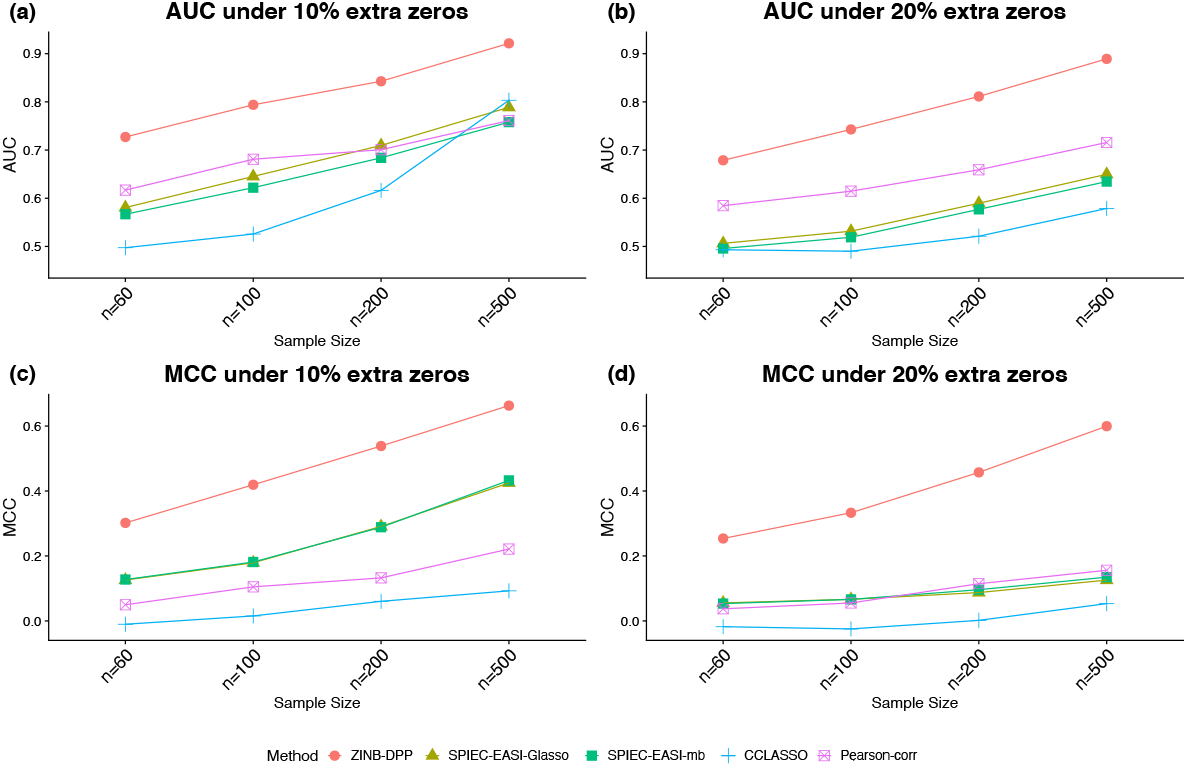
Simulated data: (a) and (b) area under the ROC curves (AUCs) and (c) and (d) area under the precision-recall curves (AUPRs) achieved by different methods under the number of taxa *p* = 40 and different sample sizes and zero proportions, averaged over 50 replicates.

**Figure 2:**
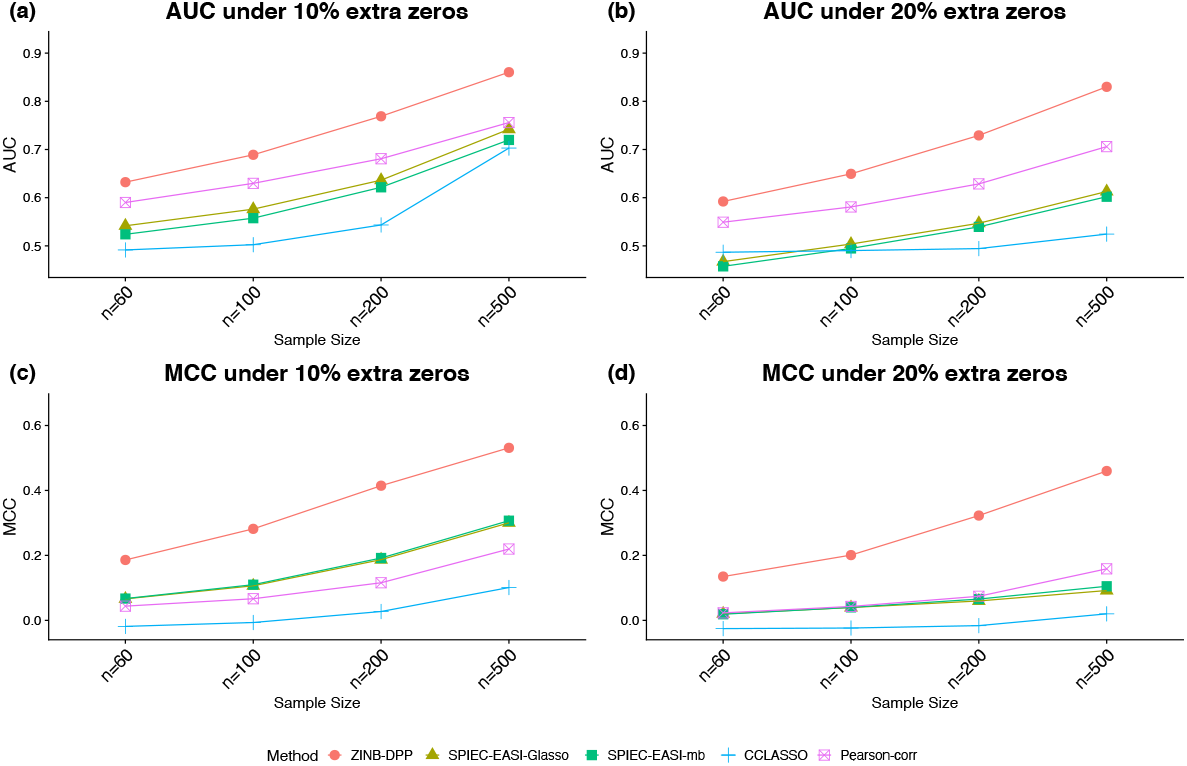
Simulated data: (a) and (b) area under the ROC curves (AUCs) and (c) and (d) area under the precision-recall curves (AUPRs) achieved by different methods under the number of taxa *p* = 60 and different sample sizes and zero proportions, averaged over 50 replicates.

Figure 3 demonstrates that our model outperformed all others on the synthetic datasets. The performances in terms of AUC under different scenarios are summarized in Figure 3(a) and (b), while those in terms of AUPR are displayed in (c) and (d). As we can see, either increasing the sample size *n* or decreasing the number of features *p* would improve the performance of all methods and lead to greater disparity between partial and pairwise correlation-based methods. In general, our HARMONIES maintained the best in all simulation and evaluation settings except for one case, where the SPIEC-EASI-mb only showed a marginal advantage (see *n* = 60 in Figure 3(c)). Interestingly, our observation confirmed a finding mentioned by Kurtz et al. (2015), that is, the SPIEC-EASI-mb was slightly better than SPIEC-EASI-Glasso in terms of AUPR under the optimal choice of the tuning parameter. As for the two correlation-based methods, we found that Pearson-corr outperformed CClasso in most of the scenarios.

**Figure 3:**
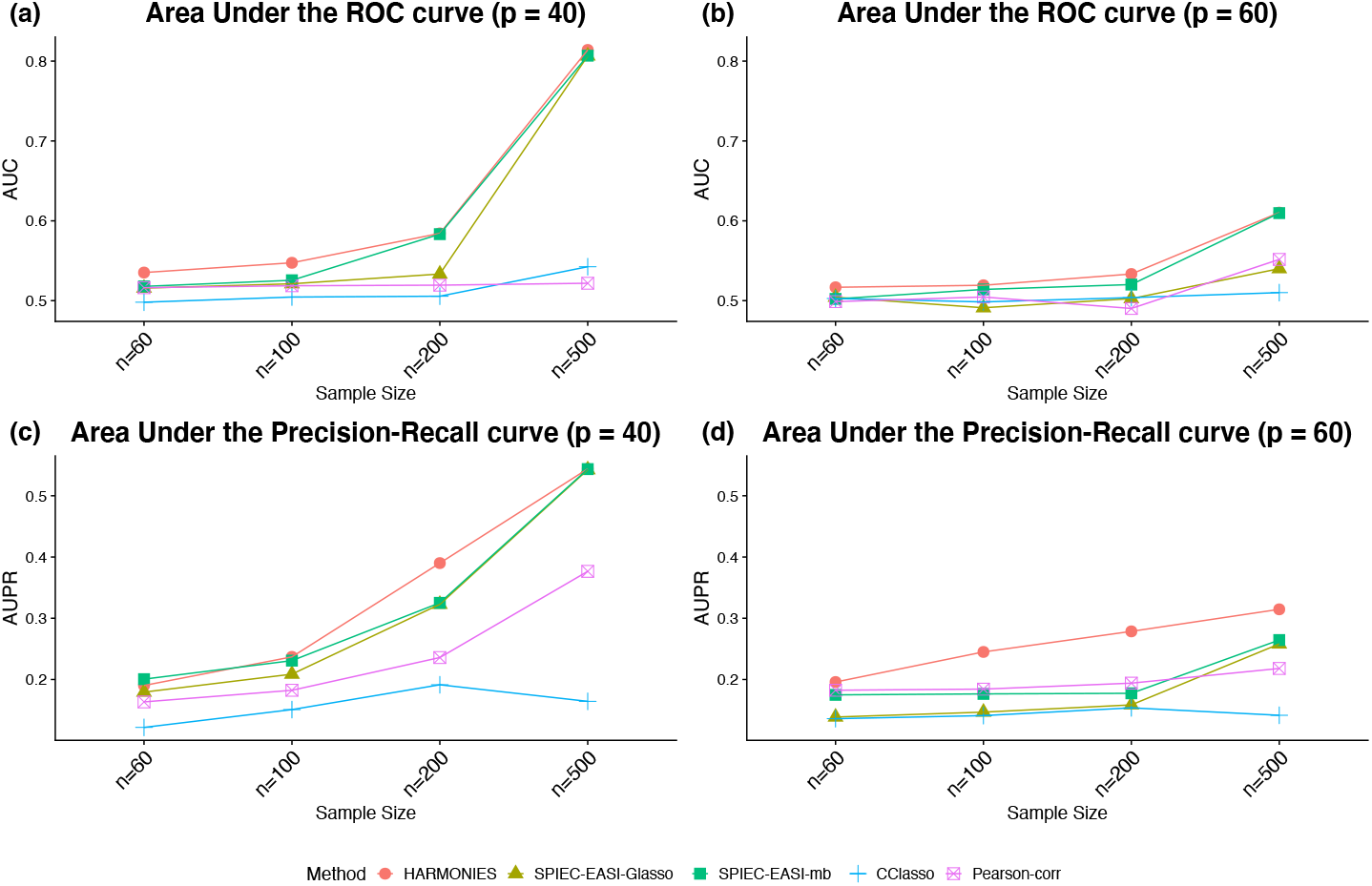
Synthetic data: (a) and (b) area under the ROC curves (AUCs) and (c) and (d) area under the precision-recall curves (AUPRs) achieved by different methods under different sample sizes and taxa numbers, averaged over 50 replicates.

### 3.2 Analysis of microbiome data from colorectal cancer patients

Colorectal cancer (CRC) is the third most common cancer diagnosed in both men and women in the United States Arnold et al. (2017). Increasing evidence from the recent studies highlights a vital role for the intestinal microbiota in malignant gastrointestinal diseases including CRC Sears and Garrett (2014); Louis et al. (2014); Drewes et al. (2016). In particular, studies have reported that dysbiosis of specific microbiota is directly associated with CRC Marchesi et al. (2011); Kostic et al. (2013); Flynn et al. (2016). The current microbiome research interests have gone beyond the discovery of disease-related microbiota, with a growing number of studies investigating the interactive associations among the microbial taxa. Using the proposed model, we interrogated the microbiome profiling data of a CRC study to determine the microbiome network structures.

We analyzed the gut microbiome dataset of a CRC study published by Feng et al. (2015). We extracted from the original cohort^1^ the 43 CRC patients and the 58 healthy controls. The original sequencing data at the genus level were quantified using curatedMetagenomicData Pasolli et al. (2017). We had *p* = 187 genera for both the 43 CRC patients and the 58 healthy controls. We implemented the HARMONIES as follows. For the CRC group, we first applied the ZINB model to obtain the normalized abundance matrix ***A***, utilizing the specifications detailed in Section 2.1. We then took the logarithmic transformation of the normalized abundance and imputed the missing values. Before implementing the proposed method, we filtered out the low abundant genera with zeros occurring more than half samples. Removing low abundant taxa is a common step in microbiome research (see e.g. Zeller et al., 2014; Qin et al., 2014; Kurtz et al., 2015; Kostic et al., 2015; Wadsworth et al., 2017; Yilmaz et al., 2019). The rationale being that these “zero-abundant” taxa may be less important in a network, which was also confirmed by our simulation study. This filtering process left 51 and 36 genera in the CRC and control group, respectively. The result from using a more relaxed filtering threshold is available in the supplement, where we kept the genera that had at least 10% nonzero observations across the samples.

Figure 4 (a) and (b) display the estimated networks for the CRC and the control group, respectively. Each node, corresponding to a genus, was named after its phylum level. All the genera shown in Figure 4 belong to six phyla in total. By using their phylum name to further categorize these distinct genera, we aimed at exploring interesting patterns among them at a higher taxonomic level. Figure S1 displays the same network using the actual genus name on each node. The node sizes are proportional to its normalized abundances in the logarithmic scale. The green or red edge indicates a positive or a negative partial correlation, respectively. And the width of an edge is proportional to the absolute value of the partial correlation coefficient. To make a clear comparison, we intentionally kept the nodes and their positions to be consistent between the two subfigures. In either of the two groups, we included a node in the current plot if there exists an edge between it with any nodes in at least one group. In general, the two groups share several edges with the same direction of partial correlations, but the majority of edges are unique within each group.

**Figure 4:**
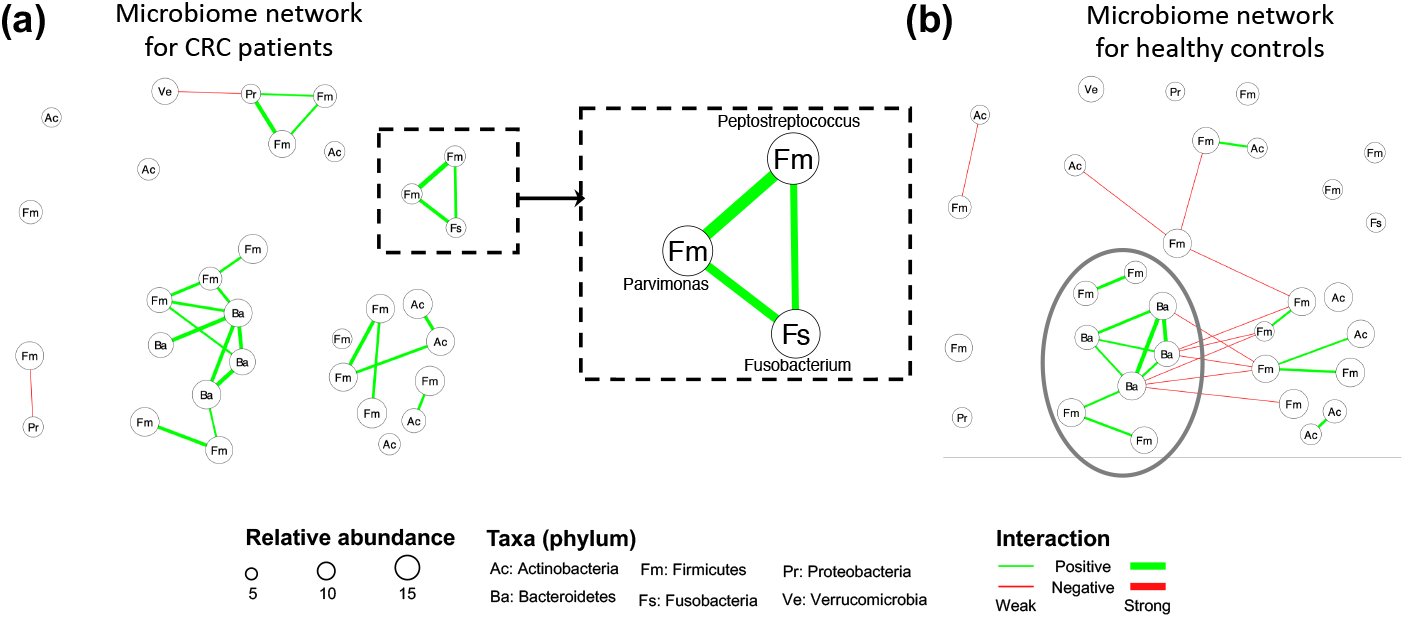
CRC case study: The estimated networks by HARMONIES for (a) CRC patients and (b) healthy controls. Increased abundances of species under the three genera (*Fusobacterium*, *Peptostreptococcus*, *Parvimonas*) in the dashed rectangular box in (a) were reported to be associated with the disease. CRC patients and healthy controls shared a similar subnetwork (composed of eight genera) circled in (b). Each node here represents a genus labeled by its phylum name. The version with distinct genus names is available in Figure S1 in the supplement.

Network estimation of the CRC group demonstrated several microbial communities. For example, three genera: *Fusobacterium*, *Peptostreptococcus*, and *Parvimonas* consisted of a unique subnetwork as highlighted in Figure 4(a). These three genera were isolated in the control group’s network, as shown in Figure 4 (b). Interestingly, specific species under these three genera have been reported as enriched taxa in CRC and related to worse clinical outcome Mima et al. (2016); Yu et al. (2017); Long et al. (2019). A previous CRC study by Kostic et al. (2013) supported the causal role of species *Fusobacterium nucleatum* (*F. nucleatum*) by showing that *F. nucleatum* promotes tumor progression by increasing both tumor multiplicity and tumor-infiltrating myeloid cells in a preclinical CRC model. Further, a recent study Long et al. (2019) demonstrated that *Peptostreptococcus anaerobius* (*P. anaerobius*) accelerated colorectal tumorigenesis in a murine CRC model. This study suggested that *P. anaerobius* directly interacted with colonic epithelial cells and also promoted CRC by modifying the tumor immune microenvironment. While the causal role of the species *Parvimonas micra* (*P. micra*) has not been biologically validated, multiple clinical studies reported an elevated level of *P. micra* in CRC patients Yu et al. (2017); Purcell et al. (2017); Dai et al. (2018). Of interest, *Parvimonas* were closely associated with animal-based diets, which have previously been shown to be significantly associated with increased risk for CRC Chan et al. (2011). The previous studies only investigated those CRC-related taxa individually, whereas a novel finding by HARMONIES analysis suggested that all the three genera were co-aggregating in CRC patients as their pairwise associations are all positive. Interestingly, in a prior study direct positive associations between *Fusobacterium* and *Peptostreptococcus*, as well as *Peptostreptococcus* and *Parvimonas*, were identified (Hibberd et al., 2017). However, there was no direct association between *Fusobacterium* and *Parvimonas*. Similarly, another study Drewes et al. (2017) found a direct co-occurrence pattern between two species: *F. nucleatum* and *P. micra*. Using HARMONIES, we could jointly identify the relationship among each pair of the three genera, conditional on all other genera. This novel subcommunity of three CRC-enriched genera formulated a recurring module, and may function as a cooperative group in CRC patients. A closer investigation of their co-occurrence pattern could potentially elucidate both their contributions to CRC and the basic biology under their relationships. Two additional novel taxa interactions were identified by HARMONIES analysis: *Streptococcus* and *Veillonella*, and *Streptococcus* and *Haemophilus*. In fact, previous CRC studies showed enrichment of these three genera or their species in CRC patients (see e.g. Geng et al., 2014; Ugai et al., 2014; Kumar et al., 2017; Koliarakis et al., 2019), but had not detected these novel interactions. In conclusion, HARMONIES may reveal how multiple CRC-related taxa could potentially promote disease progression together.

Having shared edges between the two networks suggests that the HARMONIES is robust to the edge selection. We observed that the shared edges tended to appear for those more abundant genera. For example, we circled eight genera in Figure 4 (b), and the HARMONIES suggested multiple positive partial correlations among them. For these eight genera, we observed six shared edges between the CRC and healthy control networks. Notice that all the shared edges were consistent in the association directions, and they also corresponded to the relatively stronger association in both networks (wider in the edge width). We found these shared edges tend to connect those more abundant genera (node with larger size). Indeed, the eight genera considered here belong to phyla *Bacteroidetes* and *Firmicutes*, both were in the top three most abundant phyla for CRC patients and healthy controls reported by Mori et al. (2018); Gao et al. (2015). Therefore, it was more likely that the highly abundant genera shared similar association patterns between the two groups, and the HARMONIES demonstrated its robustness by preserving these relatively stronger partial correlations among these genera. On the other hand, the network of the control group contained more negative partial correlations as shown in Figure 4 (b). Furthermore, the two edges linked to *Streptococcus* were different from the CRC group. Here, *Streptococcus* had a negative association with *Subdoligranulum* and a positive association with *Rothia*. There has been no evidence suggesting these two genera are CRC-related. Hence a further investigation is merited. Additionally, the CRC group has another distinct small subnetwork formed by the four genera, two from *Firmicutes*, one from *Proteobacteria*, and one from *Verrucomicrobia*. These group-specific associations were never reported. Lastly, we observed several interesting patterns between the two groups when summarizing the genera to their phylum levels. Genera in Firmicutes (labeled as “Fm” in Figure 4) showed more positive associations in the case group than in the control group, whereas negative associations between Firmicutes and Bacteroidetes (labeled as “Ba” in Figure 4) were more common in the control group. Again, these novel patterns still need further biological validations to elucidate their functions.

## 4 Discussion

With the advent of next-generation sequencing technology, microbiome research now has the opportunity to explore microbial community structure and to characterize the microbial ecological association for different populations or physiology conditions Kurtz et al. (2015). In this paper, we introduce HARMONIES as a statistical framework to infer sparse networks using microbiome sequencing data. It models the original count data by a zero-inflated negative binomial distribution to capture the large amount for zeros and over-dispersion, and it further implements Dirichlet process priors to account for sample heterogeneity. In contrast, current methods for microbiome network analyses rely on the compositional data, which could cause information loss due to ignoring the unique characteristics of the microbiome sequencing count data. Following the data normalization step, the HARMONIES explores the direct connections in the network by estimating the partial correlations. The results from the simulation study have demonstrated the advantage of the HARMONIES over alternative approaches under various conditions. When applied to an actual microbiome dataset, the HARMONIES suggests all the nodes be taxa at the same taxonomic level, such as species, genus, family, etc. This ensures proper biologically interpretations of those detected associations. When applied to a real CRC study, the HARMONIES revealed an intriguing community among three CRC-enriched genera. Further, shared patterns between the CRC and the control networks suggest a common community pattern of disease neutral genera. Additional studies validating the biological relevance of these microbial associations, however, will need to be conducted. Both the simulated and synthetic data showed that a larger sample size improved the performance of all the network learning methods. In practice, many disease-related microbiome studies, especially those studying rare diseases, always have small sample sizes. This limitation directly affects the estimation of the normalized matrix ***A*** from the ZINB model. Notice that for a taxon *j*, a small sample size could result in a large variance in the posterior distribution of log ***α_·j_***. However, many disease studies include reference groups where the measurements on the same taxonomic features are available. The additional information from the subjects in the reference group can potentially help improve the posterior inference of the normalized abundances. We generalized the proposed ZINB model to handle two groups, with the goal of borrowing information between groups in estimating the normalized abundances. These detailed model formula and implementation were included in the supplement (see Supplement: Infer the normalized abundances for multiple groups).

Our hybrid approach for microbiome network inference can be extended. One future direction is to incorporate the differential network analysis into the existing framework. It jointly considers the association strengths between each pair of taxa from different groups, and it compares the estimated individual networks to capture the significantly different connectivities. Our current method can infer the normalized abundances for two groups, and we provided the details steps in the supplement. However, an integrated differential network can be expected to better study the differential microbial community structure and link the communities to human health status.

## Acknowledgments

We thank Jiwoong Kim for the helpful discussions.

## Supplementary Materiel

## 1 MCMC algorithm

We start by writing the likelihood for each sample *i, i* = 1,…,*n* as

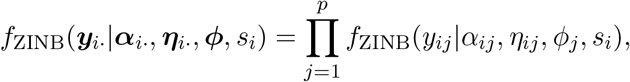

where

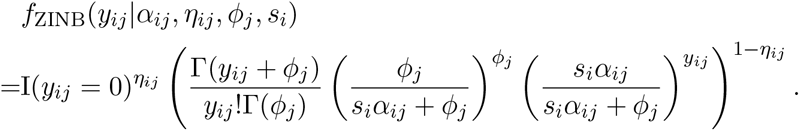

### Update of zero-inflation indicator *η_ij_*

We update each *η_ij_*, *i* = 1,…,*n, j* = 1,…,*p* that corresponds to *y_ij_* = 0 by sampling from the normalized version of the following conditional:

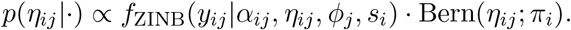

After the Metropolis-Hasting steps for all *η_ij_*, we use a Gibbs sampler to update each *π_i_*, *i* = 1,…,*n*:

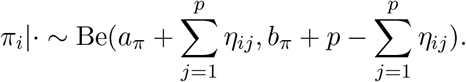

### Update of dispersion parameter *ϕ_j_*

We update each *ϕ_j_*, *j* = 1,…,*p* by using a random walk Metropolis-Hastings algorithm. We first propose a new *ϕ*_*j*_^*^ from Ga(*ϕ*_*j*_^2^/*τ_ϕ_*, *ϕ_j_*/*τ_ϕ_*) and then accept the proposed value *ϕ*_*j*_^*^ with probability min(1*, m*_MH_), where

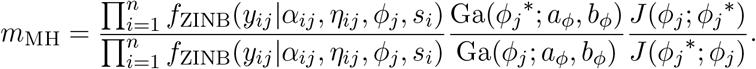

Here we use *J* (·|·) to denote the proposal probability distribution for the selected move. Note that the last term, which is the proposal density ratio, can be canceled out for this random walk Metropolis update.

### Update of size factor *s_i_*

We can rewrite Equation (2) in the main text, i.e.

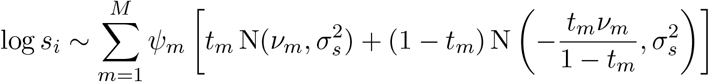

by introducing latent auxiliary variables to specify how each sample (in terms of log *s_i_*) is assigned to any of the inner and outer mixture components. More specifically, we can introduce an *n* × 1 vector of assignment indicators ***g***, with *g_i_* = *m* indicating that log *s_i_* is a sample from the *m*-th component of the outer mixture. The weight *ψ_m_* determines the probability of each value *g_i_* = *m*, with *m* = 1,…,*M*. Similarly, we can consider an *n* × 1 vector ***ϵ*** of binary elements *ϵ_i_*, where *ϵ_i_* = 1 indicates that, given *g_i_* = *m*, log *s_i_* is drawn from the first component of the inner mixture, i.e. 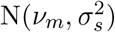 with probability *t_m_*, and *ϵ_i_* = 0 indicates that log *s_i_* is drawn from the second component of the inner mixture, i.e. 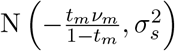, with probability 1 − *t_m_*.

Thus, the Dirichlet process prior (DPP) model can be rewritten as

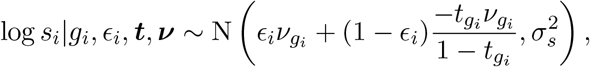

where ***t*** and ***ν*** denote the collections of *t_m_* and *ν_m_*, respectively. Therefore, the update of the size factor *s_i_*, *i* = 1,…,*n* can proceed by using a random walk Metropolis-Hastings algorithm. We propose a new 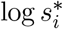 from 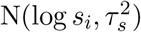 and accept it with probability min(1, m_MH_), where

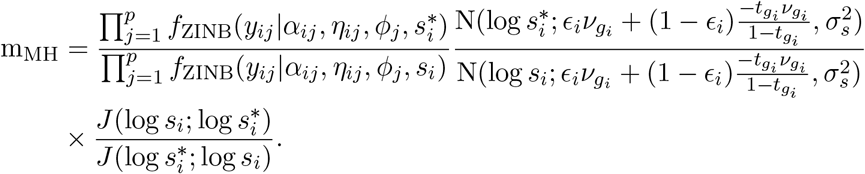

Note that the last term, which is the proposal density ratio, equals 1 for this random walk Metropolis update. Since ***g***, ***ϵ***, ***t***, and ***ν*** have conjugate full conditionals, we use Gibbs samplers to update them one after another:

- Gibbs sampler for updating *g_i_, i* = 1,…,*n*, by sampling from the normalized version of the following conditional:

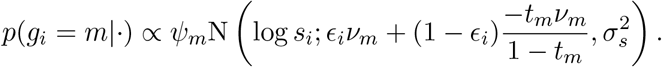
- Gibbs sampler for updating *ϵ_i_*, *i* = 1,…,*n*, by sampling from the normalized version of the following conditional:

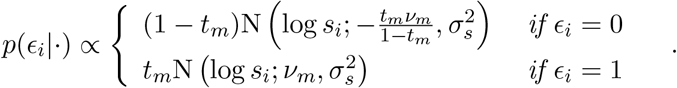
- Gibbs sampler for updating *t_m_*, *m* = 1,…,*M* :

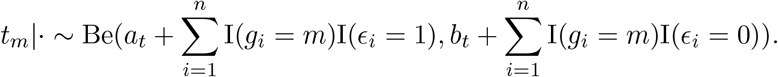
- Gibbs sampler for updating *ν_m_*, *m* = 1,…,*M* :

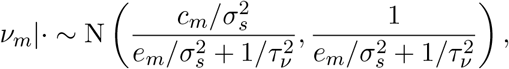

where 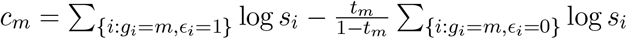 and 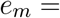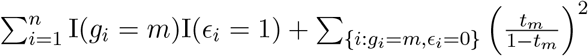.
- Gibbs sampler for updating *ψ_m_*, *m* = 1,…,*M* by stick-breaking process:

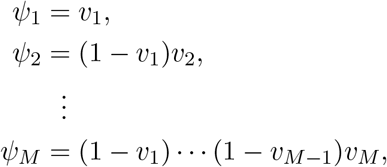

where 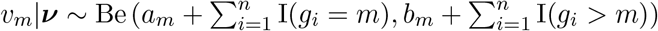.

For the sake of convenience, we have copied Equation (3) in the main text here,

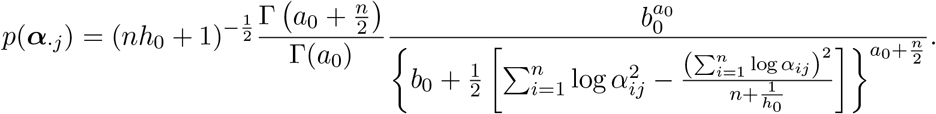

### Update of normalized abundance

We update each *α_ij_*, *i* = 1,…,*n, j* = 1,…,*p* by using a Metropolis-Hastings random walk algorithm. We first propose a new *α*_*ij*_^*^ from 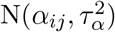, and then accept the proposed value with probability min(1, m_MH_), where

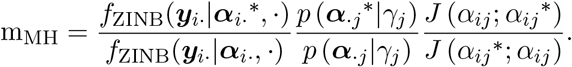

Note that the last term, which is the proposal density ratio, equals 1 for this random walk Metropolis update.

#### Supplement to Section 3.2: Analysis of microbiome data from colorectal cancer patients

We carried out a simple sensitivity analysis to evaluate the model performance to the choice of the filtering threshold. As discussed in Section 3.2, we filtered out the genera with more than 50% of nonzero counts across the samples. Here, we changed the threshold from 50% to 10%. The new threshold left 92 and 84 genera in the CRC and control group, respectively. Figure S2 shows the network inferred under this new setting.

Our first observation is that there were a large number of similarities between the networks. For instance, in the CRC group, the relatively stronger associations remained the same, such as positive associations between *Bacteroides* and *Alistipes*, *Barnesiella* and *Alistipes*, *Blautia* and *Dorea*, *Streptococcus* and *Haemophilus*. Though the new CRC network did not show any negative partial correlations (denoted by the red edges), the negative associations in the original network were relatively weak and might not be as stable as the other edges. Notably, the “positive triangle” among the three genera *Fusobacterium*, *Peptostreptococcus*, and *Parvimonas* was again confirmed here. In the control group, the strong associations can be still detected, such as the positive associations between *Alistipes* and *Parabacteroides*, *Alistipes* and *Bacteroides*, and negative associations between *Bacteroides* and *Dorea*. It is interesting to point out that the networks based on the new threshold tended to have fewer edges.

We also observed that increasing the number of genera in the networks could introduce novel associations. For example, genera *Pantoea* and *Escherichia* were dropped from the original CRC group network, yet they established a positive association in the new network. Similarly, genera *Pantoea*, *Escherichia*, *Abiotrophia*, *Oxalobacter*, and *Desulfovibrio* were of low abundance in the healthy controls and hence were not considered in the model before. However, they were connected in the new network. These results estimated from those highly sparse genera may hint further biological validations.

## 2 Infer the normalized abundances for multiple groups

In practice, when there are two groups of subjects in a microbiome study (e.g., subjects with two distinct phenotypes), the sequencing data usually include measurements on the same taxonomic features for all the subjects. Then, if the abundance of a taxon *j* does not differ between two groups, we can improve the posterior influence of log ***α***_·*j*_ by merging two groups to increase the sample size. On the other hand, if the taxon is associated with subject’s condition, i.e., a taxon that changes its abundance between two groups in the study, the inference of log ***α***_·*j*_ should rely on each subject group.

With the goal of borrowing information to improve the posterior inference for certain taxa, we combine the original count matrix from two different groups, to generate the count matrix ***Y***_*n*×*p*_. Here, the sample size is *n* = *n*_1_ + *n*_2_, with *n*_1_*, n*_2_ representing the number of subjects in the first and the second groups, respectively. Meanwhile, we let ***z*** = (*z*_1_,…,*z_n_*)^*T*^ to allocate the *n* subjects into two groups, with *z_i_* = 1 or 2 indicating the group label of subject *i*. In practice, if taxon *j* is irrelevant to the subject’s phenotype, its abundances should not be differentiating between two groups. However, if taxon *j* is associated with the disease, its abundance could either increase or decrease from healthy subjects to patients. Therefore, we model the normalized abundance *α_ij_* as following:

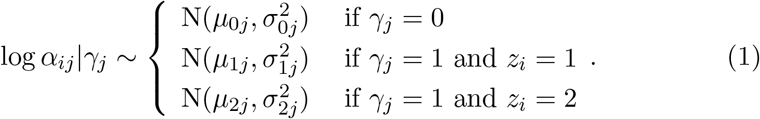

Here, *γ_j_* is a latent binary variable, with *γ_j_* = 1 if taxon *j* is differentially abundant between two groups, and *γ_j_* = 0 otherwise. For the taxa with *γ_j_* = 0, we can borrow information between groups to increase the sample size in estimating the corresponding posterior of log ***α***_·*j*_. As an extension to Section 2.1 where we assume 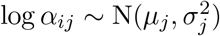, the current model includes *μ*_0*j*_, *μ*_1*j*_, and *μ*_2*j*_ as the mean parameters for the normal mixture model. Again, we take the conjugate Bayesian approach and impose the following priors for the parameters in the normal mixture model: 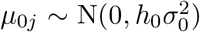, 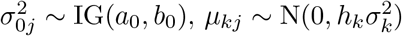 and 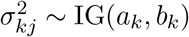 for *k* = 1, 2.

The estimation of *γ_j_*’s determines the resulted normalized abundance matrix. Specifically, for taxon *j* with *γ_j_* = 0, we can impute the zeros due to missing by the posterior mean of log ***α***_·*j*_ calculated using information from both groups. As an extension to Equation (3) in the main text, the posterior of *α_·j_ |γ_j_* is as following:

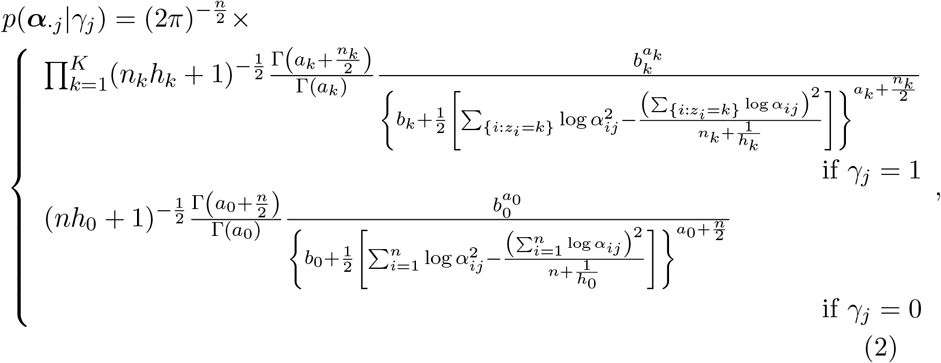

Therefore, we can obtain the posterior mean of log ***α***_·*j*_ by averaging over the log-transformed MCMC samples of ***α***_·*j*_ after burn-in.

---

**Figure 1:**
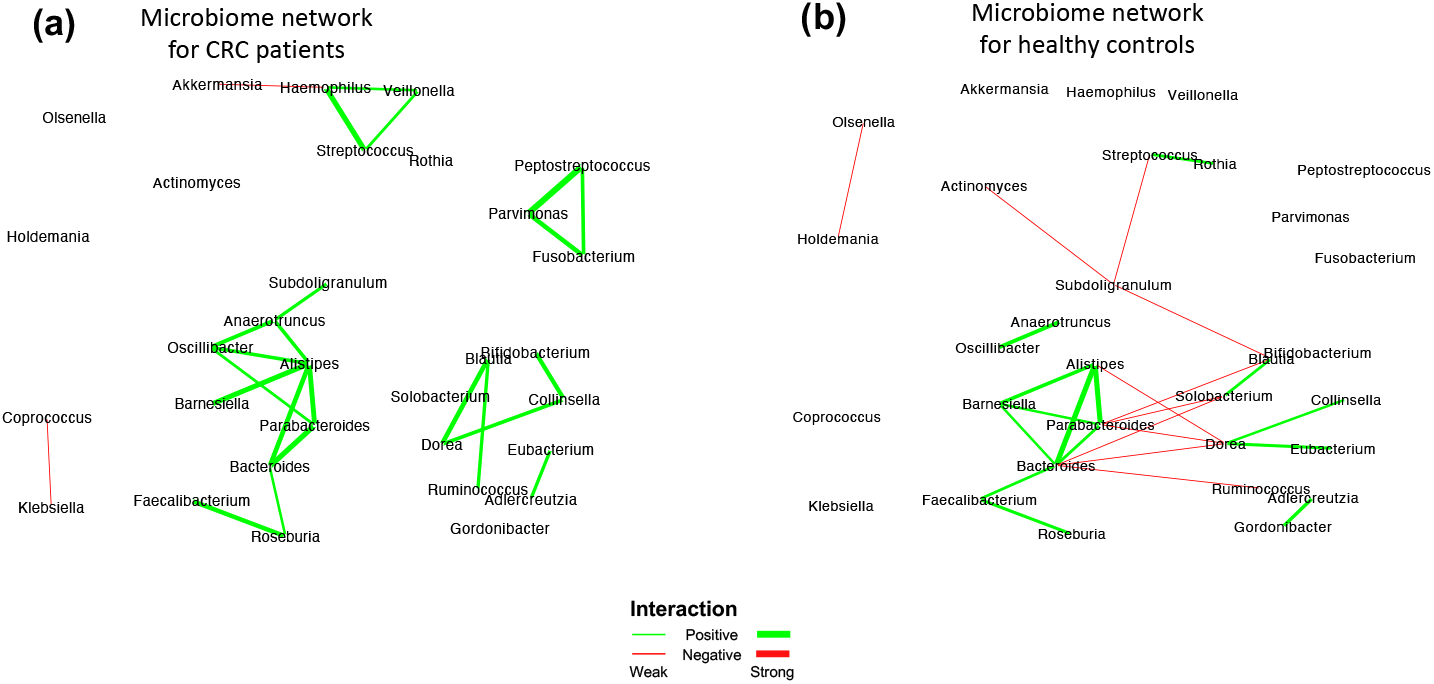
CRC case study: The estimated networks by HARMONIES for (a) CRC patients and (b) healthy controls. All nodes are labeled in their genus names.

**Figure 2:**
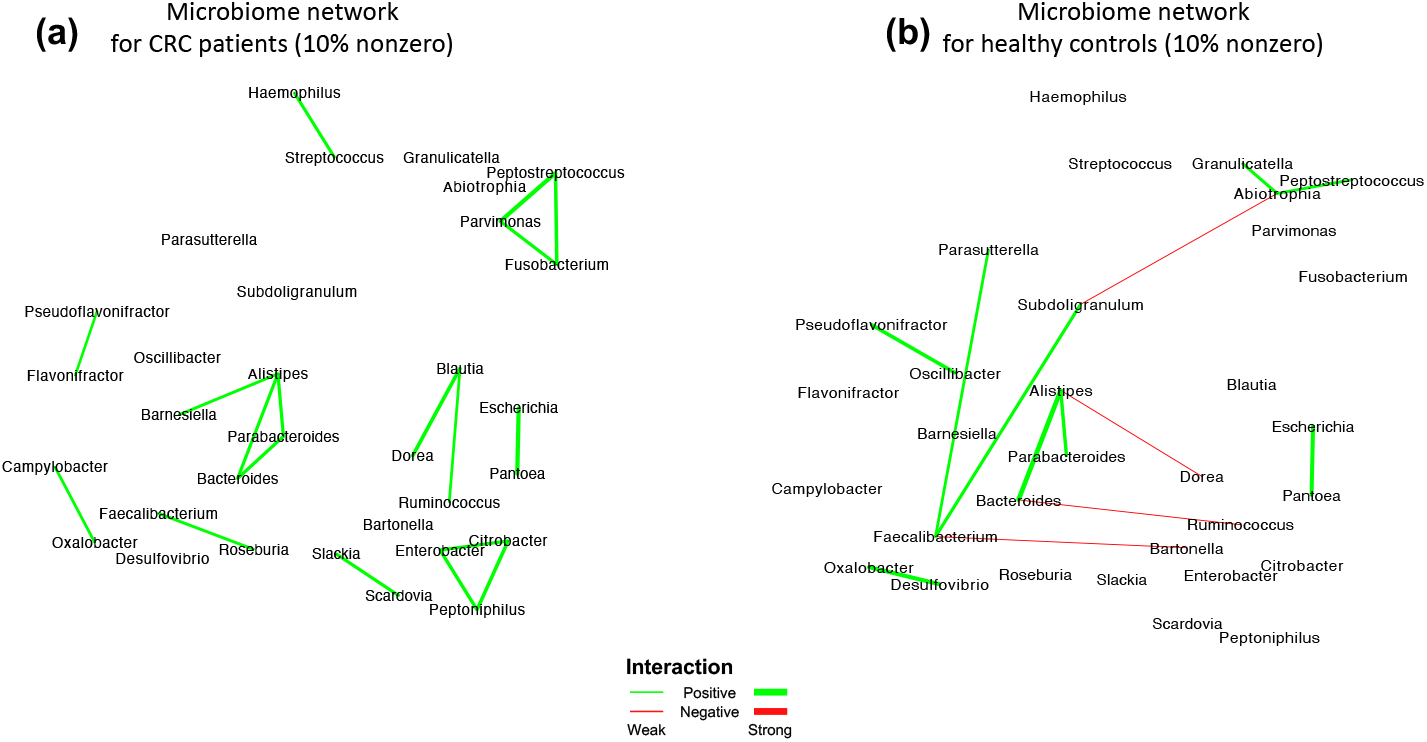
CRC case study: The estimated networks by HARMONIES for (a) CRC patients and (b) healthy controls. The nodes are genera that have at least 10% nonzero observations across the samples in each group.

## Footnotes

1 The original metagenomic shotgun sequencing data from the fecal samples are available in the European Bioinformatics Institute Database (accession number ERP008729)

## References

Anders, S. and Huber, W. (2010). Differential expression analysis for sequence count data. Genome Biology 11, R106

Arnold, M., Sierra, M. S., Laversanne, M., Soerjomataram, I., Jemal, A., and Bray, F. (2017). Global patterns and trends in colorectal cancer incidence and mortality. Gut 66, 683–691

Ban, Y., An, L., and Jiang, H. (2015). Investigating microbial co-occurrence patterns based on metagenomic compositional data. Bioinformatics 31, 3322–3329

Bullard, J. H., Purdom, E., Hansen, K. D., and Dudoit, S. (2010). Evaluation of statistical methods for normalization and differential expression in mRNA-Seq experiments. BMC Bioinformatics 11, 94

Cario, M. C. and Nelson, B. L. (1997). Modeling and generating random vectors with arbitrary marginal distributions and correlation matrix. Tech. rep., Citeseer

Chan, D. S., Lau, R., Aune, D., Vieira, R., Greenwood, D. C., Kampman, E., et al. (2011). Red and processed meat and colorectal cancer incidence: meta-analysis of prospective studies. PloS One 6, e20456

Dai, Z., Coker, O. O., Nakatsu, G., Wu, W. K., Zhao, L., Chen, Z., et al. (2018). Multi-cohort analysis of colorectal cancer metagenome identified altered bacteria across populations and universal bacterial markers. Microbiome 6, 70

Drewes, J. L., Housseau, F., and Sears, C. L. (2016). Sporadic colorectal cancer: microbial contributors to disease prevention, development and therapy. British Journal of Cancer 115, 273

Drewes, J. L., White, J. R., Dejea, C. M., Fathi, P., Iyadorai, T., Vadivelu, J., et al. (2017). High-resolution bacterial 16S rRNA gene profile meta-analysis and biofilm status reveal common colorectal cancer consortia. NPJ Biofilms and Microbiomes 3, 1–12

Fang, H., Huang, C., Zhao, H., and Deng, M. (2015). CClasso: correlation inference for compositional data through lasso. Bioinformatics 31, 3172–3180

Faust, K. and Raes, J. (2016). CoNet app: inference of biological association networks using Cytoscape. F1000Research 5

Feng, Q., Liang, S., Jia, H., Stadlmayr, A., Tang, L., Lan, Z., et al. (2015). Gut microbiome development along the colorectal adenoma-carcinoma sequence. Nature Communications 6, 6528

Flynn, K. J., Baxter, N. T., and Schloss, P. D. (2016). Metabolic and community synergy of oral bacteria in colorectal cancer. Msphere 1

Friedman, J., Hastie, T., and Tibshirani, R. (2008). Sparse inverse covariance estimation with the graphical lasso. Biostatistics 9, 432–441

Gao, Z., Guo, B., Gao, R., Zhu, Q., and Qin, H. (2015). Microbiota disbiosis is associated with colorectal cancer. Frontiers in Microbiology 6, 20

Geng, J., Song, Q., Tang, X., Liang, X., Fan, H., Peng, H., et al. (2014). Co-occurrence of driver and passenger bacteria in human colorectal cancer. Gut Pathogens 6, 26

Gevers, D., Kugathasan, S., Denson, L. A., Vázquez-Baeza, Y., Van Treuren, W., Ren, B., et al. (2014). The treatment-naive microbiome in new-onset Crohn’s disease. Cell Host & Microbe 15, 382–392

Hibberd, A. A., Lyra, A., Ouwehand, A. C., Rolny, P., Lindegren, H., Cedgåard, L., et al. (2017). Intestinal microbiota is altered in patients with colon cancer and modified by probiotic intervention. BMJ Open Gastroenterology 4, e000145

Koliarakis, I., Messaritakis, I., Nikolouzakis, T. K., Hamilos, G., Souglakos, J., and Tsiaoussis, J. (2019). Oral bacteria and intestinal dysbiosis in colorectal cancer. International Journal of Molecular Sciences 20, 4146

Kostic, A. D., Chun, E., Robertson, L., Glickman, J. N., Gallini, C. A., Michaud, M., et al. (2013). *Fusobacterium nucleatum* potentiates intestinal tumorigenesis and modulates the tumor-immune microenvironment. Cell Host & Microbe 14, 207–215

Kostic, A. D., Gevers, D., Siljander, H., Vatanen, T., Hyötyläinen, T., Hämäläinen, A.-M., et al. (2015). The dynamics of the human infant gut microbiome in development and in progression toward type 1 diabetes. Cell Host & Microbe 17, 260–273

Kumar, R., Herold, J. L., Schady, D., Davis, J., Kopetz, S., Martinez-Moczygemba, M., et al. (2017). *Streptococcus gallolyticus* subsp. *gallolyticus* promotes colorectal tumor development. PLoS Pathogens 13

Kurtz, Z. D., Müller, C. L., Miraldi, E. R., Littman, D. R., Blaser, M. J., and Bonneau, R. A. (2015). Sparse and compositionally robust inference of microbial ecological networks. PLoS Computational Biology 11, e1004226

Kyung, M., Gill, J., and Casella, G. (2011). Sampling schemes for generalized linear Dirichlet process random effects models. Statistical Methods & Applications 20, 259–290

Lam, C. and Fan, J. (2009). Sparsistency and rates of convergence in large covariance matrix estimation. Annals of Statistics 37, 4254

Lee, J. and Sison-Mangus, M. (2018). A bayesian semiparametric regression model for joint analysis of microbiome data. Frontiers in Microbiology 9, 522

Li, Q., Cassese, A., Guindani, M., and Vannucci, M. (2019). Bayesian negative binomial mixture regression models for the analysis of sequence count and methylation data. Biometrics 75, 183–192

Li, Q., Guindani, M., Reich, B. J., Bondell, H. D., and Vannucci, M. (2017). A Bayesian mixture model for clustering and selection of feature occurrence rates under mean constraints. Stat. Anal. Data Min. 10, 393–409

Liu, H., Roeder, K., and Wasserman, L. (2010). Stability approach to regularization selection (StARS) for high dimensional graphical models. In Advances in Neural Information Processing Systems. 1432–1440

Lo, C. and Marculescu, R. (2017). MPLasso: Inferring microbial association networks using prior microbial knowledge. PLoS Computational Biology 13, e1005915

Long, X., Wong, C. C., Tong, L., Chu, E. S., Szeto, C. H., Go, M. Y., et al. (2019). Peptostreptococcus anaerobius promotes colorectal carcinogenesis and modulates tumour immunity. Nature Microbiology, 1–12

Louis, P., Hold, G. L., and Flint, H. J. (2014). The gut microbiota, bacterial metabolites and colorectal cancer. Nature Reviews Microbiology 12, 661–672

Marchesi, J. R., Dutilh, B. E., Hall, N., Peters, W. H., Roelofs, R., Boleij, A., et al. (2011). Towards the human colorectal cancer microbiome. PloS One 6, e20447

Meinshausen, N., Bühlmann, P., et al. (2006). High-dimensional graphs and variable selection with the lasso. The Annals of Statistics 34, 1436–1462

Menéndez, P., Kourmpetis, Y. A., ter Braak, C. J., and van Eeuwijk, F. A. (2010). Gene regulatory networks from multifactorial perturbations using Graphical Lasso: application to the DREAM4 challenge. PloS One 5, e14147

Metzker, M. L. (2010). Sequencing technologies—the next generation. Nature Reviews Genetics 11, 31

Mima, K., Nishihara, R., Qian, Z. R., Cao, Y., Sukawa, Y., Nowak, J. A., et al. (2016). *Fusobacterium nucleatum* in colorectal carcinoma tissue and patient prognosis. Gut 65, 1973–1980

Mori, G., Rampelli, S., Orena, B. S., Rengucci, C., De Maio, G., Barbieri, G., et al. (2018). Shifts of faecal microbiota during sporadic colorectal carcinogenesis. Scientific Reports 8

Newton, M. A., Noueiry, A., Sarkar, D., and Ahlquist, P. (2004). Detecting differential gene expression with a semiparametric hierarchical mixture method. Biostatistics 5, 155–176

Oh, J. H. and Deasy, J. O. (2014). Inference of radio-responsive gene regulatory networks using the graphical lasso algorithm. BMC Bioinformatics 15, S5

Pasolli, E., Schiffer, L., Manghi, P., Renson, A., Obenchain, V., Truong, D. T., et al. (2017). Accessible, curated metagenomic data through experimenthub. Nature Methods 14, 1023

Paulson, J. N., Stine, O. C., Bravo, H. C., and Pop, M. (2013). Differential abundance analysis for microbial marker-gene surveys. Nature Methods 10, 1200

Peng, J., Wang, P., Zhou, N., and Zhu, J. (2009). Partial correlation estimation by joint sparse regression models. Journal of the American Statistical Association 104, 735–746

Purcell, R. V., Visnovska, M., Biggs, P. J., Schmeier, S., and Frizelle, F. A. (2017). Distinct gut microbiome patterns associate with consensus molecular subtypes of colorectal cancer. Scientific Reports 7, 11590

Qin, N., Yang, F., Li, A., Prifti, E., Chen, Y., Shao, L., et al. (2014). Alterations of the human gut microbiome in liver cirrhosis. Nature 513, 59–64

Robinson, M. D. and Oshlack, A. (2010). A scaling normalization method for differential expression analysis of RNA-seq data. Genome Biology 11, R25

Sears, C. L. and Garrett, W. S. (2014). Microbes, microbiota, and colon cancer. Cell Host & Microbe 15, 317–328

Taddy, M. A., Kottas, A., et al. (2012). Mixture modeling for marked Poisson processes. Bayesian Analysis 7, 335–362

Tipton, L., Müller, C. L., Kurtz, Z. D., Huang, L., Kleerup, E., Morris, A., et al. (2018). Fungi stabilize connectivity in the lung and skin microbial ecosystems. Microbiome 6, 12

Ugai, T., Norizuki, M., Mikawa, T., Ohji, G., and Yaegashi, M. (2014). Necrotizing fasciitis caused by haemophilus influenzae type b in a patient with rectal cancer treated with combined bevacizumab and chemotherapy: a case report. BMC Infectious Diseases 14, 198

Wadsworth, W. D., Argiento, R., Guindani, M., Galloway-Pena, J., Shelburne, S. A., and Vannucci, M. (2017). An integrative bayesian dirichlet-multinomial regression model for the analysis of taxonomic abundances in microbiome data. BMC Bioinformatics 18, 94

Weiss, S., Van Treuren, W., Lozupone, C., Faust, K., Friedman, J., Deng, Y., et al. (2016). Correlation detection strategies in microbial data sets vary widely in sensitivity and precision. The ISME journal 10, 1669

Yilmaz, B., Juillerat, P., Øyås, O., Ramon, C., Bravo, F. D., Franc, Y., et al. (2019). Microbial network disturbances in relapsing refractory Crohn’s disease. Nature Medicine 25, 323–336

Yu, J., Feng, Q., Wong, S. H., Zhang, D., yi Liang, Q., Qin, Y., et al. (2017). Metagenomic analysis of faecal microbiome as a tool towards targeted non-invasive biomarkers for colorectal cancer. Gut 66, 70–78

Zeller, G., Tap, J., Voigt, A. Y., Sunagawa, S., Kultima, J. R., Costea, P. I., et al. (2014). Potential of fecal microbiota for early-stage detection of colorectal cancer. Molecular Systems Biology 10

Zhao, H. and Duan, Z.-H. (2019). Cancer Genetic Network Inference Using Gaussian Graphical models. Bioinformatics and Biology Insights 13, 1177932219839402

